# Aging and interferon gamma response drive the phenotype of neutrophils in the inflamed joint

**DOI:** 10.1101/2021.11.16.465905

**Authors:** Ricardo Grieshaber-Bouyer, Tarik Exner, Nicolaj S. Hackert, Felix A. Radtke, Scott A. Jelinsky, Olha Halyabar, Alexandra Wactor, Elham Karimizadeh, Jorge Schettini, A. Helena Jonsson, Deepak A. Rao, Lauren A. Henderson, Carsten Müller-Tidow, Hanns-Martin Lorenz, Guido Wabnitz, James A. Lederer, Angela Hadjipanayis, Peter A. Nigrovic

**Author notes:** **Correspondence** to R.G-B and P.A.N.: Ricardo Grieshaber-Bouyer, MD, Division of Rheumatology, Department of Medicine V (Hematology, Oncology and Rheumatology), Heidelberg University Hospital, Im Neuenheimer Feld 410, 69120 Heidelberg, Germany, Peter A. Nigrovic, MD, Chief, Division of Immunology, Boston Children’s Hospital, Karp Family Research Building, Room 10211, One Blackfan Circle, Boston, MA 02115, USA.

## Abstract

**Objectives:** Neutrophils are typically the most abundant leukocyte in arthritic synovial fluid. We sought to understand changes that occur in neutrophils as they migrate from blood to joint.

**Methods:** We performed RNA sequencing of neutrophils from healthy human blood, arthritic blood, and arthritic synovial fluid, comparing transcriptional signatures with those from murine K/BxN serum transfer arthritis. We employed mass cytometry to quantify protein expression and sought to reproduce the synovial fluid phenotype *ex vivo* in cultured healthy blood neutrophils.

**Results:** Blood neutrophils from healthy donors and patients with active arthritis exhibited largely similar transcriptional signatures. By contrast, synovial fluid neutrophils exhibited more than 1,600 differentially expressed genes. Gene signatures identified a prominent response to interferon gamma (IFNγ), as well as to tumor necrosis factor, interleukin 6, and hypoxia, in both humans and mice. Mass cytometry also found healthy and arthritic donor blood neutrophils largely indistinguishable but revealed a range of neutrophil phenotypes in synovial fluid defined by downregulation of CXCR1 and upregulation of FcγRI, HLA-DR, PD-L1, ICAM-1 and CXCR4. Reproduction of key elements of this signature in cultured blood neutrophils required both IFNγ and prolonged culture.

**Conclusions:** Circulating neutrophils from arthritis patients resemble those from healthy controls, but joint fluid cells exhibit a network of changes, conserved across species, that implicate IFNγ response and aging as complementary drivers of the synovial neutrophil phenotype.

**KEY MESSAGES:** *What is already known about this subject?:*

- Neutrophils are central in the effector phase of inflammatory arthritis but their phenotypic heterogeneity in inflamed synovial fluid is poorly understood.

*What does this study add?:*

- RNA-seq and mass cytometry identify a hallmark phenotype of neutrophils in synovial fluid consisting of upregulated ICAM-1, HLA-DR, PD-L1, Fc receptors and CXCR4.
- Transcriptomics highlight an IFNγ response signature conserved across humans and mice.
- In vitro experiments implicate aging and IFNγ as complementary factors orchestrating the synovial fluid neutrophil phenotype.

*How might this impact on clinical practice or future developments?:*

- Understanding the specific features of neutrophils in the arthritic joint may disclose opportunities for safe therapeutic targeting of this lineage.

## INTRODUCTION

Inflammatory arthritis encompasses a broad spectrum of diseases affecting adults and children[1]. The pathogenesis of non-infectious arthritis is correspondingly varied, with upstream mechanisms that include autoantibodies, T cells, autoinflammatory mechanisms, and crystals[2]. Despite this remarkable pathogenic diversity, a ubiquitous feature of arthritic joint fluid is an abundance of neutrophils, a canonical innate immune effector cell required for immune defense but also for many forms of pathogenic inflammation.

Compelling evidence confirms that neutrophils are key pathogenic contributors in arthritis. Neutrophils from human joints exhibit altered surface markers and function consistent with activation[3-7]. Synovial neutrophils elaborate pro-inflammatory factors such as interleukin (IL)-1, leukotriene B4, citrullinated peptides, and neutrophil extracellular traps[8-11]. Neutrophils activated by adherent immune complexes degrade articular cartilage[12]. Finally, mice with defects specific to the neutrophil compartment – for example, depleted of neutrophils, congenitally deficient in neutrophils, with neutrophils lacking key effector molecules, or subject to neutrophil migratory blockade – exhibit dense resistance to experimental arthritis[11, 13-17]. Therefore, understanding the phenotype of synovial fluid neutrophils is essential to understanding the biology of arthritis and may reveal novel opportunities for therapeutic intervention[18].

Advances in cellular characterization offer new ways to understand neutrophils. Transcriptomic analysis provides a hypothesis-independent examination of the activity of cells at the gene expression level, informing the relationships between populations of cells. For example, studies using single-cell RNA sequencing (scRNAseq) recently established that murine neutrophils represent a single lineage, differentiating along a developmental continuum termed ‘neutrotime’, rather than a branched network of committed subtypes[19]. Indeed, aging is well recognized as a modulator of neutrophil phenotype and function[20, 21]. Importantly, however, the relationship between a neutrophil’s transcriptome and its surface protein signature varies markedly with context[19]. For example, neutrophil activation is accompanied by rapid mobilization of the surface integrin CD11b from an intracellular pool and cleavage-mediated loss of the surface selectin CD62L[22]. Mass cytometry (cytometry by time of flight, CyTOF) allows simultaneous determination of dozens of surface and intracellular markers in each cell, albeit restricted by investigator choice as to the markers most likely to prove informative[23].

To understand the changes that occur in neutrophils as they enter the inflamed joint, we applied low-input RNAseq to purified neutrophils, sorting cells by a known dichotomous surface marker of undetermined function, CD177, to eliminate potential confounding by variation in the CD177^pos^ neutrophil fraction within the population[24]. We compared human neutrophils with scRNAseq transcriptome data from murine neutrophils, both circulating and from autoantibody-mediated neutrophil-driven K/BxN serum transfer arthritis[25]. We employed CyTOF to define markers of neutrophil differentiation and function, followed by confirmatory *in vitro* studies using flow cytometry. We establish that blood neutrophils from healthy and arthritic donors are largely similar but find that synovial fluid neutrophils differ markedly from blood neutrophils in a manner that implicates both interferon gamma (IFNγ) response and cell aging in the resulting phenotype.

## METHODS

Demographic characteristics and materials and methods are available in the supplementary materials.

## RESULTS

### Transcriptomic characterization of circulating neutrophils

We performed low-input RNAseq on blood neutrophils from 15 healthy and 16 arthritic donors (**Supplementary Table 1**), calculating Pearson correlation coefficients between samples based on the expression of all genes. Hierarchical clustering of these correlation coefficients revealed no separation by disease state, suggesting little divergence of transcriptional phenotype (**Figure 1A**). At an FDR of 0.05 (indicated by red line), blood neutrophils from healthy and arthritic donors differed in only three genes: *DNAJB9* (DnaJ Heat Shock Protein Family (Hsp40) Member B9), *DDIT4* (DNA damage inducible transcript 4) and *SCO2* (Synthesis of Cytochrome C Oxidase 2), all modestly upregulated in arthritis (**Figure 1B, C**).

**Figure 1.**
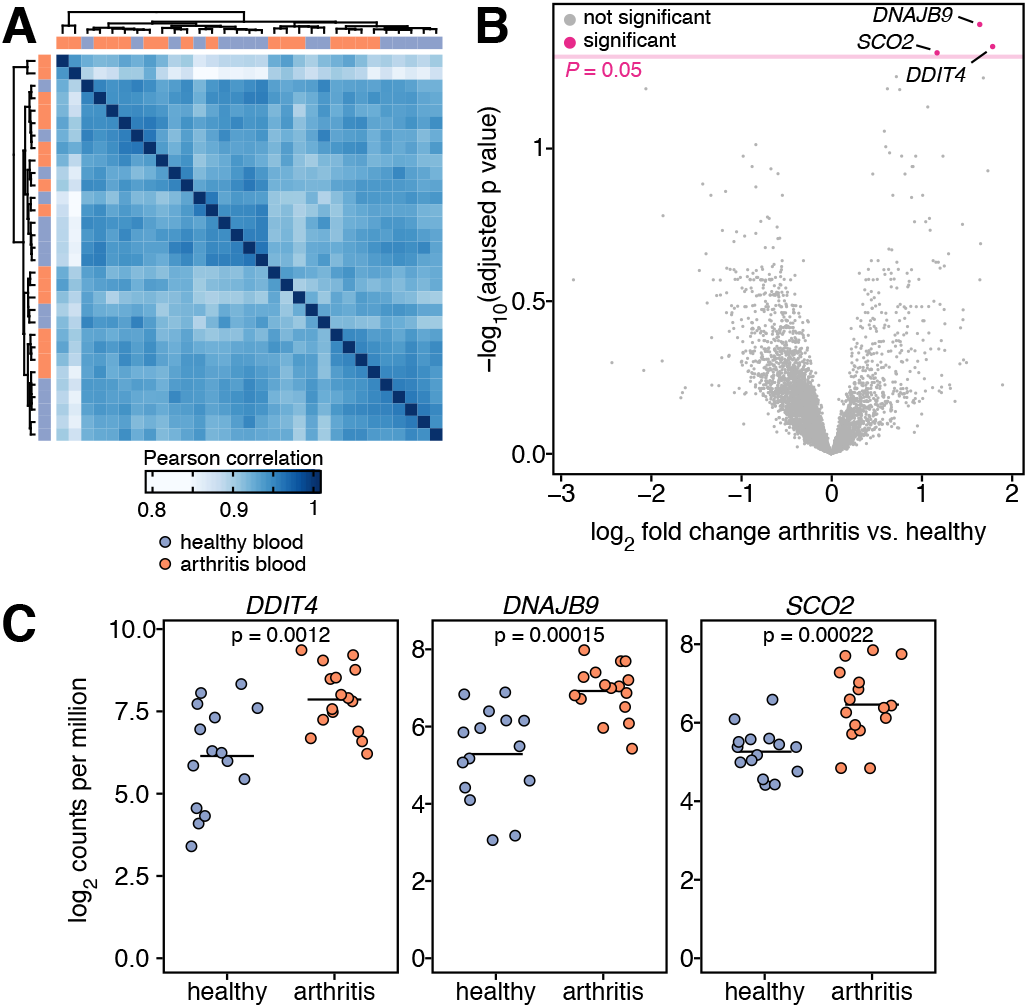
Transcriptomic similarity of blood neutrophils from healthy controls and patients with inflammatory arthritis. **(A)** Hierarchical clustering of Pearson correlation coefficients between individual blood samples based on the expression of all genes reveals complete overlap between the two groups. **(B)** Volcano plot of differentially expressed genes (at FDR 0.05) between blood neutrophils from healthy and arthritic donors. **(C)** *DNAJB9, DDIT4* and *SCO2* are overexpressed in blood of inflammatory arthritis patients compared to healthy controls. N = 15 healthy controls, N = 16 arthritis patients.

### Synovial fluid neutrophils display an IFNγ response

We performed the same analysis in 16 paired contemporaneous peripheral blood and synovial fluid samples from patients with active arthritis requiring therapeutic joint aspiration. Hierarchical clustering revealed separation into two groups as a function of location (**Figure 2A**). At │log fold change│≥ 1 and FDR 0.05, 1,657 genes were differentially expressed, of which 939 were downregulated and 718 upregulated (**Figure 2B**).

**Figure 2.**
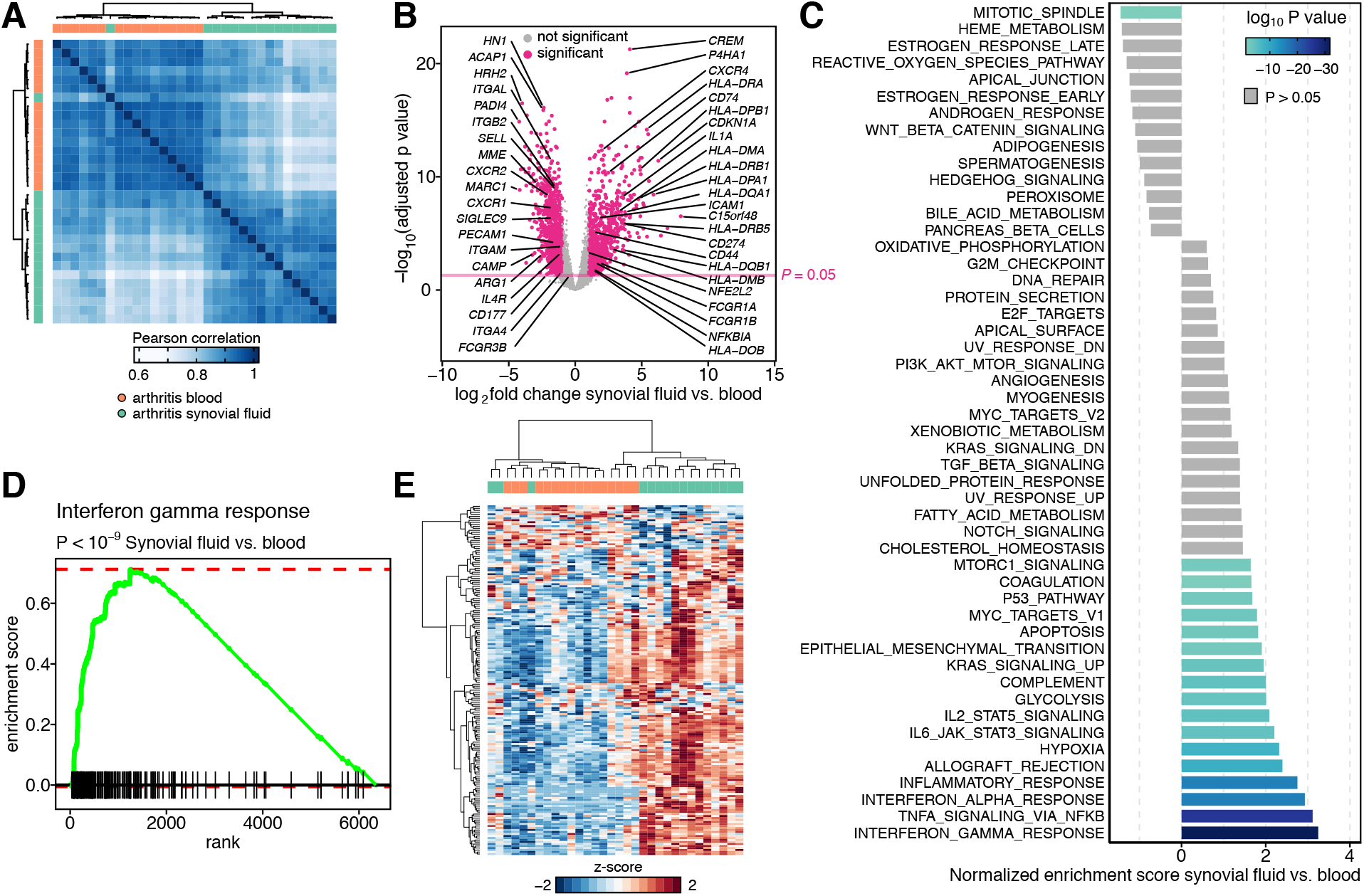
Synovial fluid neutrophils are enriched for IFNγ response genes. **(A)** Hierarchical clustering of Pearson correlation coefficients between paired peripheral blood and synovial fluid samples based on the expression of all genes shows strong separation based on tissue. **(B)** 1,657/6,350 genes are differentially expressed at log_2_ fold change ≥ 1 and FDR of 0.05 between peripheral blood and synovial fluid neutrophils. **(C)** Gene Set Enrichment Analysis of differentially expressed genes in synovial fluid vs. blood neutrophils. **(D)** Enrichment plot of the IFNγ response signature in synovial fluid neutrophils. **(E)** Expression heatmap of IFNγ response genes in synovial fluid neutrophils reveals strong separation between blood and synovial fluid. N = 16 paired blood and synovial fluid inflammatory arthritis samples.

To understand these genes in terms of functional programs, we employed Gene Set Enrichment Analysis using the established 50 hallmark gene sets. Signatures of TNF and IL-6 response were easily detected, as was response to hypoxia (**Figure 2C**). However, the most prominent gene set in synovial fluid neutrophils was IFNγ response (**Figure 2C, 2D**). Analysis of genes up-regulated in response to IFNγ (HALLMARK_INTERFERON_GAMMA_RESPONSE) revealed that most (93/175) expressed IFNγ target genes were highly induced in synovial fluid neutrophils, including the class II molecules *CD74, HLA-DMA, HLA-DRB1* and *HLA-DQA1*; *CD274* (encoding PD-L1); and *FCGR1A* (encoding CD64, the high-affinity IgG receptor FcγRI) (**Figure 2E**). These observations show that phenotypic deviation of neutrophils in arthritis is primarily in the joints rather than in the circulation, at least at the transcriptional level, and suggest a prominent role for IFNγ in driving the phenotype of synovial fluid neutrophils.

### Conserved responses of human and murine neutrophils in inflammatory arthritis

Neutrophils are indispensable for onset and perpetuation of joint inflammation in mice[8-11, 14-16]. To test whether the transcriptional changes we observed in human synovial neutrophils are conserved across species, we compared our human dataset to a microarray-based transcriptional atlas of neutrophils from the blood of healthy mice and joints of mice undergoing K/BxN serum transfer arthritis[25]. We restricted the combined dataset to 5,520 one-to-one gene orthologs according to ENSEMBL version 100[26]. Of genes with orthologs significantly upregulated in human (578) and murine (226) synovial fluid neutrophils, 97 were shared across species, far more than expected by chance (95% confidence interval for chance overlap 16–34 genes as defined by random resampling 20,000 times, *P* = 2.7 × 10^−7^, **Figure 3A**). Similarly, downregulated genes across human (774) and mouse (174) neutrophils in synovial fluid shared significant overlap with 75 genes, compared to 23–41 expected by chance (*P* = 1.3 × 10^−11^, **Figure 3A**). In murine synovial fluid neutrophils, enhanced expression was observed in IFNγ target genes including *CD274* (encoding PD-L1) and the MHC class II gene *HLADQB1*; indeed an IFNγ signature was one of the key functional patterns observed, with highly skewed representation of IFNγ response genes (adjusted *P* < 0.001, **Figure 3B,C**). These findings establish that gene expression changes, including an IFNγ response signature, are shared by human and mice synovial fluid neutrophils.

**Figure 3.**
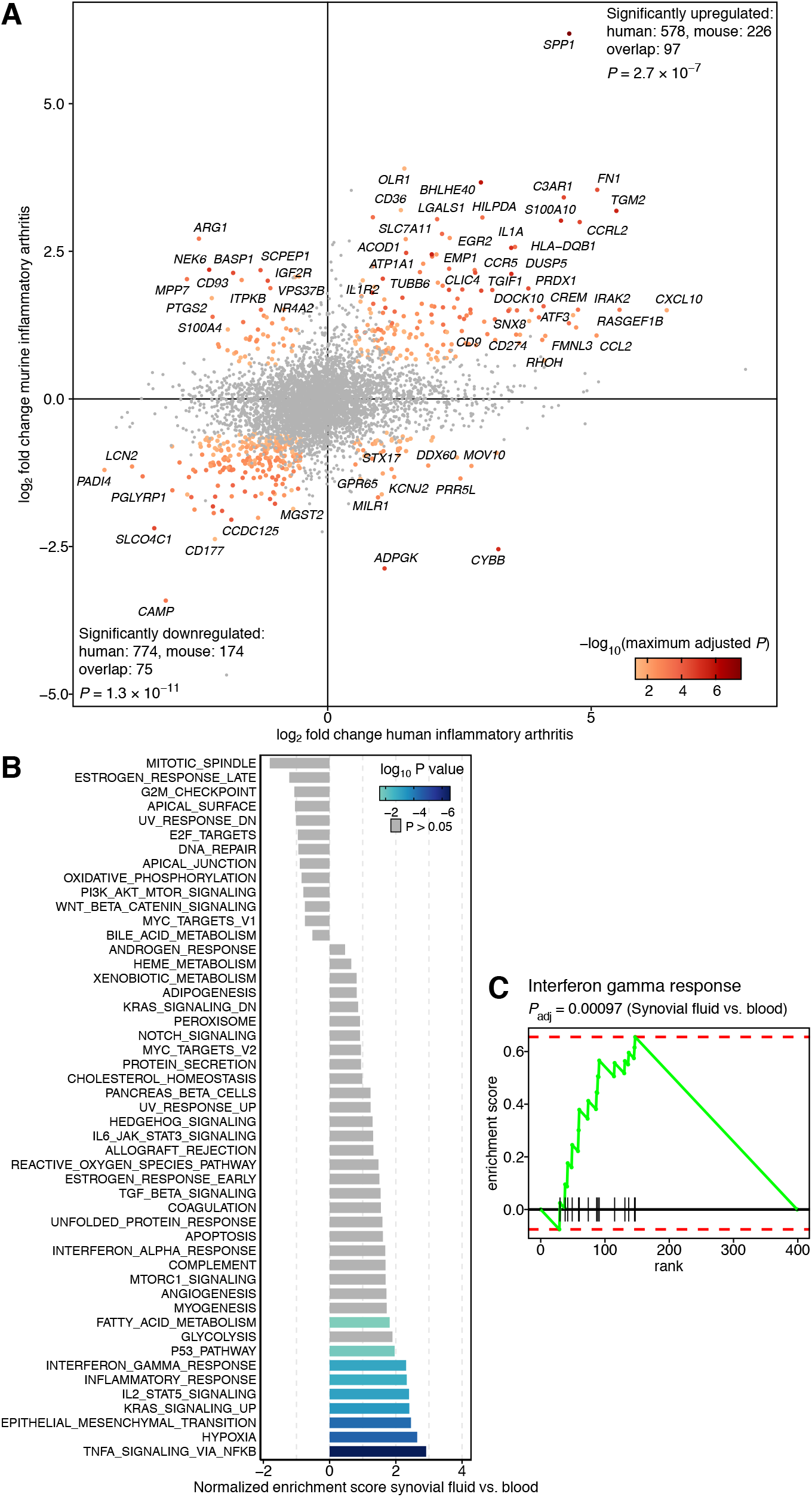
Cross species analysis of neutrophil gene expression in inflamed synovial fluid. **(A)** Depicted is the log_2_ fold change of gene expression in human (x) vs. murine (y) synovial fluid neutrophils compared to blood neutrophils. Only genes with with one-to-one orthologs are shown and genes with adjusted P < 0.05 in both comparisons and | log_2_ fold change | ≥ 0.75 are highlighted. Genes are conservatively colored by highest P value. **(B)** Significantly differentially expressed genes were ranked by log_2_ fold change and Gene Set Enrichment Analysis was performed on hallmark gene sets. **(C)** Enrichment plot of the hallmark gene set “Interferon gamma response”. Only genes with one-to-one orthologs between mice and humans are shown; for gene symbols, the human symbol is shown.

### CyTOF confirms joint-specific activation of human neutrophils in inflammatory arthritis

We created a custom CyTOF panel containing 39 human surface and intracellular markers related to neutrophil activation, chemokine receptors, antigen presentation, adhesion factors and co-stimulatory molecules (**Supplementary Methods**). CyTOF studies were performed in 33 samples (9 healthy volunteer donors, 8 blood samples from patients with inflammatory arthritis, and 16 synovial fluid samples, including 7 contemporaneous blood/synovial fluid pairs; **Supplementary Table 1**).

To analyze global data structure, we extracted median expression values for each protein in each sample and calculated Spearman correlation coefficients between samples based on expression data. Hierarchical clustering revealed complete overlap in peripheral blood neutrophils between healthy donors and patients with inflammatory arthritis, indicating few systematic differences in global protein expression (**Figure 4A**). Correspondingly, we found differential expression only of a single marker, CD64, between healthy and arthritic donor peripheral blood neutrophils after correction for multiple comparisons (**Figure 4C**). These results mirror our transcriptomic findings and show similarity of blood neutrophils between healthy and arthritic donors.

**Figure 4.**
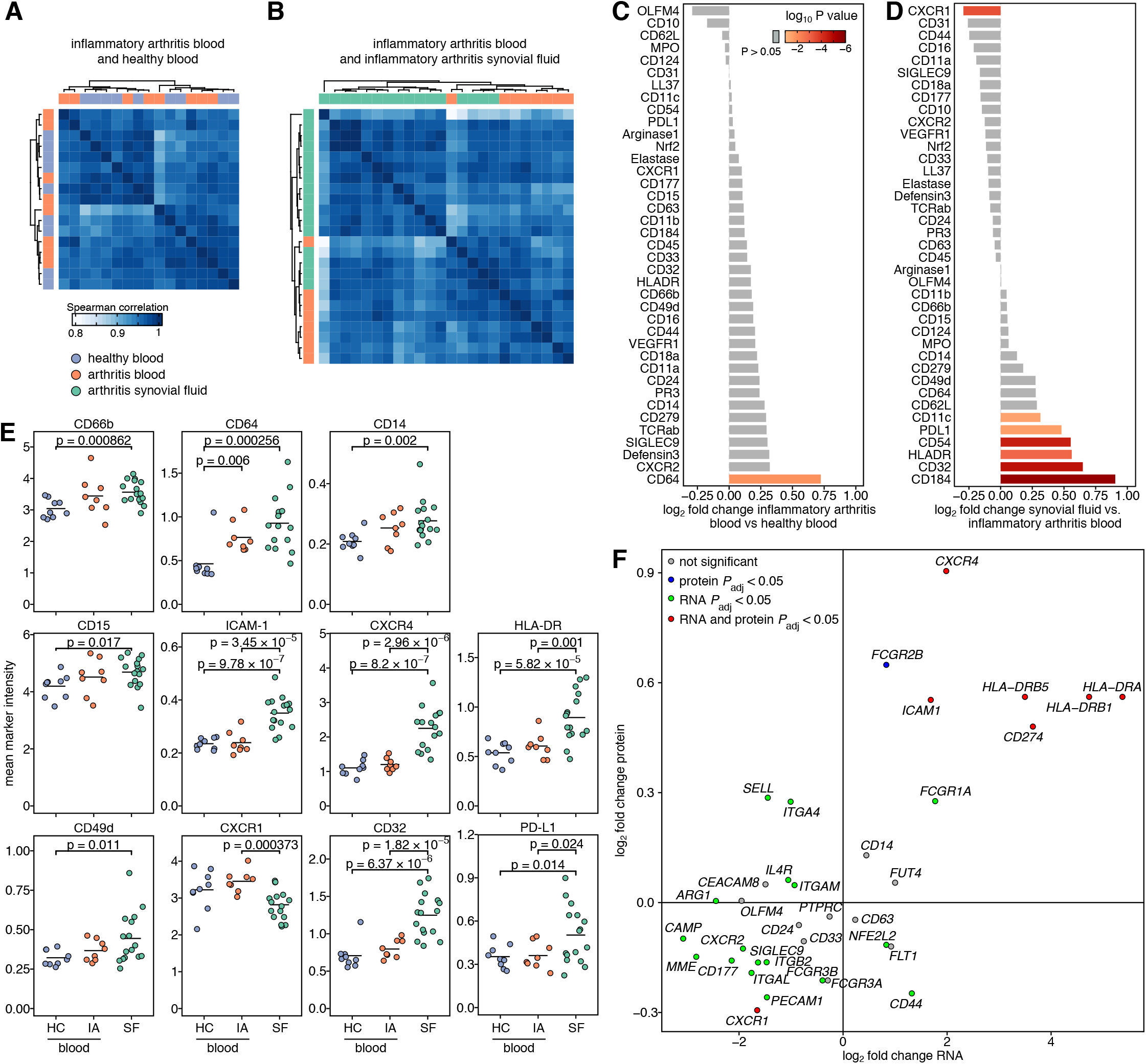
Mass cytometry analysis of neutrophils. **(A)** Hierarchical clustering of Spearman correlation coefficients between blood neutrophils from healthy donors and patients with inflammatory arthritis based on global neutrophil protein expression. **(B)** Differential expression analysis of global neutrophil marker expression in the peripheral blood. **(C)** Hierarchical clustering of Spearman correlation coefficients between peripheral blood and synovial fluid samples. **(D)** Differential expression analysis of global neutrophil marker expression between peripheral blood and synovial fluid. **(E)** Average expression of significantly differentially expressed markers per sample. **(F)** Comparison of gene- and protein expression differences between blood and synovial fluid neutrophils identifies a hallmark synovial fluid phenotype. HC = healthy control; IA = inflammatory arthritis; SF = synovial fluid

By contrast, comparison of blood and synovial fluid revealed a strong separation driven by tissue (**Figure 4B**). This separation was driven by multiple differentially expressed proteins in synovial fluid neutrophils including downregulation of CXCR1 and upregulation of the integrin CD11c, PD-L1, ICAM-1, HLA-DR, the low-affinity Fc receptor CD32 (FcγRII) and CXCR4 (CD184), the receptor for CXCL12/SDF-1 that retains neutrophils in inflamed sites [27] (**Figure 4D,E**). CD64 was also overexpressed but did not reach significance compared to arthritic donor blood due to already higher expression (**Figure 4E**). Compared to healthy blood neutrophils, synovial fluid neutrophils overexpressed CD64, the activation and lineage markers CD66b and CD15, the LPS co-receptor CD14, and the integrin CD49d (**Figure 4E**).

PD-L1, HLA-DR and CD64 are upregulated in neutrophils exposed to IFNγ, consistent with our transcriptomic signature data[28-34]. The IL-8 receptor CXCR1 was downregulated, potentially reflecting agonist-mediated internalization of this G-protein-coupled receptor. No change was noted in granule proteins, including for primary (azurophilic) granules (including MPO, proteinase 3 [PR3], arginase 1), secondary granules (including LL-37 / cathelicidin, CD177, OLFM4), and tertiary granules (arginase 1). Results for all markers are shown in **Supplementary Figure 3**.

We investigated how well expression differences in RNA and protein match each other. We found that downregulation of *CXCR1* and upregulation of *CXCR4, ICAM1, HLA-DRA, HLA-DRB1, HLA-DRB5* and *CD274* (encoding PD-L1) were highly concordant between RNA and protein (**Figure 4F**). Upregulation of *FCGR1A* (CD64) and *FCGR2B* (CD32) was also observed on both RNA and protein level but was significant only at either gene (*FCGR1A*) or protein (CD32) level. This set of genes and their protein products thus constitute hallmarks of the synovial fluid neutrophil phenotype.

### Continuous and discrete neutrophil phenotypes

To define neutrophil heterogeneity at the single-cell level, we performed Uniform Manifold Approximation and Projection (UMAP) dimensionality reduction on our CyTOF data. Initial results were dominated by the two known dichotomously expressed neutrophil proteins, CD177 and OLFM4 (**Supplementary Figure 4A**). Cells from peripheral blood and synovial fluid were evenly distributed across CD177^pos/neg^ and OLFM4^pos/neg^ populations in the UMAP embedding, suggesting that the phenotypic changes distinguishing blood and synovial fluid neutrophils operate evenly across these markers (**Supplementary Figure 4B**). Accordingly, we did not detect any significant differences in frequency of neutrophil subsets defined by CD177 or OLFM4 between blood and synovial fluid (**Supplementary Figure 4C**).

To neutralize this dominant impact, we excluded CD177, OLFM4, and the CD177-anchored enzyme PR3 from consideration and repeated dimensionality reduction. We observed a striking separation between resting blood neutrophils and synovial fluid cells, single-cell findings that mirrored our transcriptomic results (**Figure 5A**). Synovial fluid neutrophils concentrated in two primary clusters, termed here SFN1 and SFN2 and observed across individual donors (**Figure 5A** and **Supplemental Figure 5**). For analysis purposes, we forced neutrophils into k = 20 clusters, again excluding CD177, OLFM4, and PR3, with the goal of maximizing the opportunity to identify distinct phenotypic states (**Figure 5B**). Individual markers varied among the 20 clusters, confirming neutrophil heterogeneity at the single-cell level (**Figure 5C**). Examining the frequency of cells belonging to each cluster, we observed considerable divergence among donors, limiting statistical power in this relatively small sample size. Clusters 10–12 were particularly overrepresented in synovial fluid, representing the bulk of neutrophils in SFN2 (**Figure 5D**). Neutrophils in Clusters 10-11 expressed high levels of CXCR4 (CD184) and Cluster 12 cells additionally expressed the IFNγ markers HLA-DR, PD-L1 and CD64. The SFN1 population was contained within Cluster 2 and was driven primarily by a single donor, although it trended higher across multiple samples in synovial fluid vs. blood. By contrast, blood neutrophils were enriched for Clusters 1, 8, 9, and 16, with only Cluster 9 (expressing high levels of granule proteins) and Cluster 16 (expressing granule proteins, OLFM4, and CD124, the alpha chain of the IL-4 and IL-13 receptors) achieving statistical significance (**Figure 5D**).

**Figure 5.**
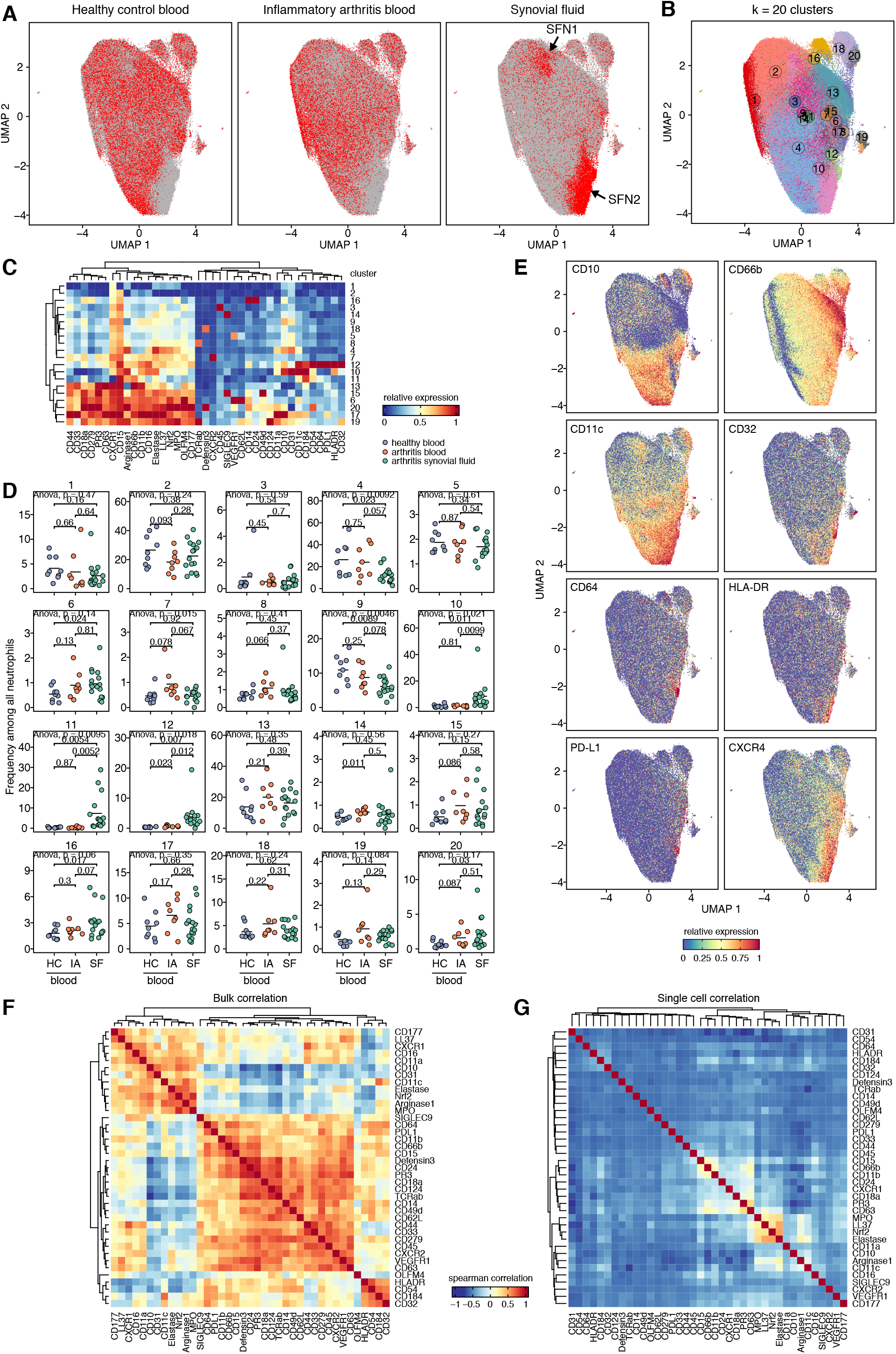
Continuous and discrete neutrophil phenotypes. **(A)** UMAP embedding of single-cell CyTOF data separates blood neutrophils and synovial fluid cells. **(B)** Overclustering of neutrophils into 20 groups captures neutrophil heterogeneity across blood and synovial fluid. **(C)** Heterogeneity in marker expression between the 20 clusters. **(D)** Change in frequency of different neutrophil phenotypes across conditions. **(E)** Gradients of marker expression characterize synovial fluid neutrophils. Correlation between markers on a per-sample **(F)** and single-cell **(G)** level identifies clusters of co-expressed markers. HC = healthy control; IA = inflammatory arthritis; SF = synovial fluid

Expression of differentially expressed markers between blood and synovial fluid revealed that most markers follow expression gradients (**Figure 5E**). Broadly, two gradients could be observed: a gradient from top to bottom that included many granule proteins and likely reflecting maturation (CD10, Nrf2, arginase 1, CD11a, elastase, LL-37, CD31, MPO, OLFM4, CD177, CD184/CXCR4) and a gradient from left to right likely reflecting activation (CD66b, CD11b, CD15, CD16, CXCR1, CD45) (**Supplementary Figure 5**). Small populations of interest included Clusters 8 and 19 (expressing TCRαβ, equally rare in blood and synovial fluid) and Cluster 6, expressing VEGFR1 and therefore potentially representing pro-angiogenic neutrophils[35], significantly increased in synovial fluid compared to healthy blood.

Notably, not all upregulated markers were expressed on the same cells. For example, CXCR4-high neutrophils expressed variable amounts of HLA-DR and PD-L1 (**Figure 5E**). Correlating expression intensity on a per-sample bulk level revealed a positive correlation between markers defining the core synovial fluid phenotype: HLA-DR, ICAM-1, CXCR4 and CD32 (**Figure 5F**). Analysis at the single-cell level confirmed a correlation between general activation markers CD11b, CD15 and CD66b and within a cluster of granule proteins (MPO, LL-37, Nrf2, Elastase) but not between CXCR4, HLA-DR and PD-L1 (**Figure 5G**).

Together, these results show that CXCR4+, HLA-DR+ and PD-L1+ neutrophils are expanded in inflamed synovial fluid and that expression of these markers peaks in different cells, confirming that the inflamed environment features divergent neutrophil phenotypes.

### IFNγ and aging drive blood neutrophils toward a synovial fluid phenotype

Since both transcriptomic and proteomic analysis revealed a strong IFNγ response signature in synovial fluid neutrophils, we hypothesized that stimulation with IFNγ could recapitulate the synovial fluid phenotype in healthy blood neutrophils. As expected, viability dropped from nearly 100% at beginning of culture to 71% after 2 days of culture at 37°C. Stimulation with IFNγ significantly extended neutrophil survival to 87% (**Figure 6A**).

**Figure 6.**
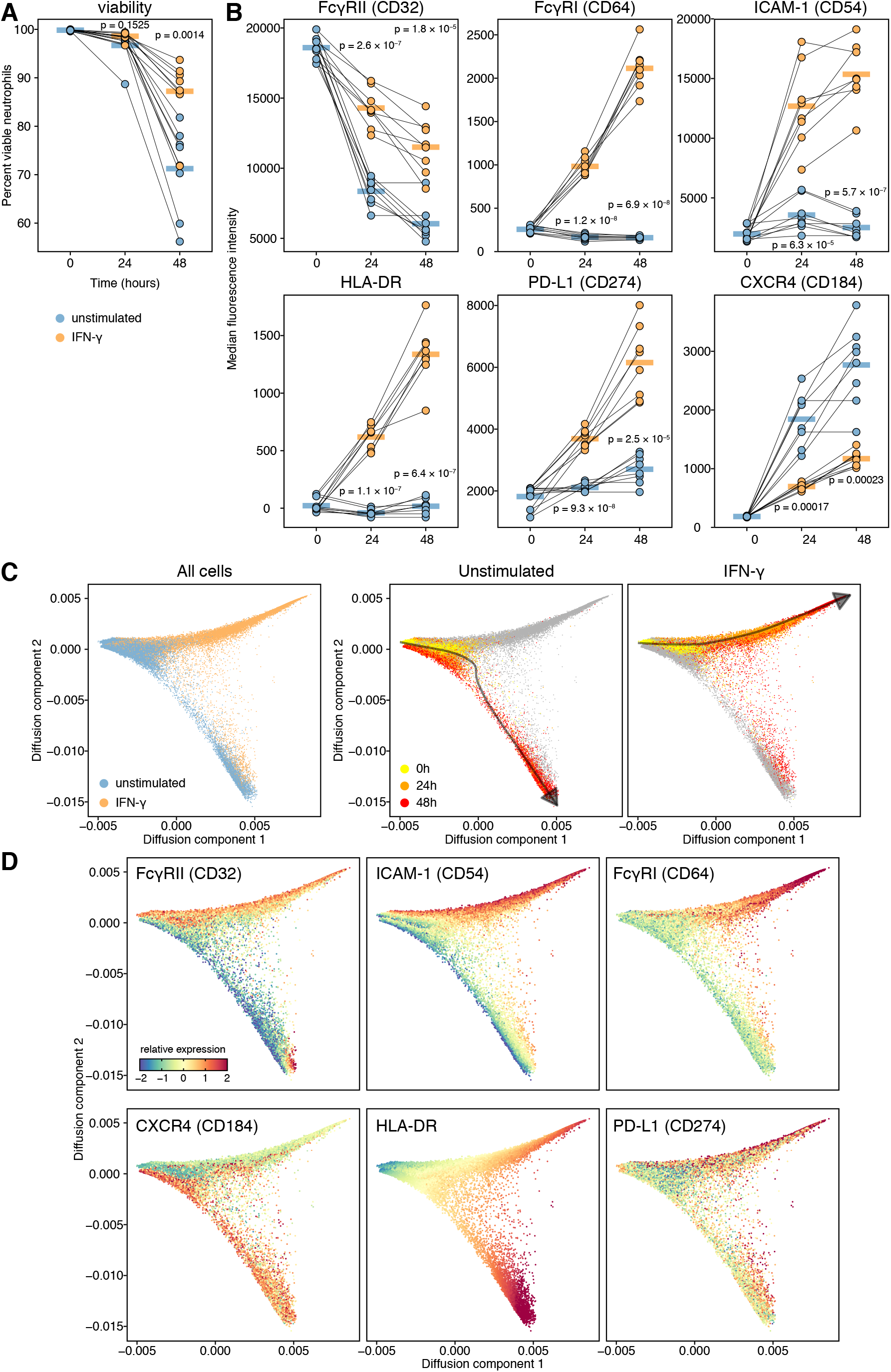
Progressive aging and response to interferon gamma recapitulate the synovial fluid phenotype in vitro. **(A)** Stimulation with IFNγ extends the lifetime of neutrophils *in vitro*. **(B)** Effect of aging and IFNγ on the expression of key surface markers. **(C)** Diffusion map of unstimulated and IFNγ stimulated neutrophils cultured over 48 hours. **(D)** Expression of key surface markers on the diffusion map.

IFNγ stimulation prevented downregulation of CD32 and significantly upregulated CD64, ICAM-1, HLA-DR and PD-L1 (**Figure 6B**). CXCR4 expression was not detectable in freshly isolated neutrophils but increased with time in culture, consistent with its known role as a marker of neutrophil aging[21]. Interestingly, CXCR4 expression was reduced by cytokine stimulation, indicating either an impact on CXCR4 expression specifically or a broader effect on the neutrophil aging program (**Figure 6B**).

Based on those findings, we hypothesized that cytokine stimulation and aging were complementary in establishing the synovial fluid neutrophil phenotype. We therefore analyzed unstimulated and stimulated neutrophils together in a single diffusion map. This analysis revealed a marked divergence in phenotypes between cells left unstimulated and those incubated with IFNγ (**Figure 6C**). CXCR4 expression was highest at the most distant pole of the unstimulated trajectory (**Figure 6D**). Conversely, IFNγ robustly upregulated HLA-DR, PD-L1 and ICAM-1 (**Figure 6D**). Thus, the combination of aging and exposure to IFNγ, but not either alone, yielded a neutrophil phenotype resembling that of synovial fluid neutrophils.

## DISCUSSION

Synovial fluid neutrophils are the hallmark of inflammatory arthritis[36]. We employed low-input RNA sequencing and CyTOF to characterize neutrophils from healthy donor blood and from blood and synovial fluid of patients with active arthritis. Whereas circulating neutrophils exhibited few changes with disease state, synovial fluid neutrophils displayed consistent phenotypic deviation implicating two conceptually orthogonal influences: response to local mediators, most prominently IFNγ, and cell aging.

The marked alteration in mRNA expressed by synovial fluid neutrophils is consistent with the growing understanding of neutrophils as highly dynamic cells that remain transcriptionally active throughout their lifespan[19, 37, 38]. This adaptability may be of particular consequence in neutrophils recruited to inflamed sites such as the arthritic joint, since cytokines can prolong neutrophil half-life from a baseline of 8-20 hours to several days[39].

Transcriptional signatures observed here included response to mediators of established importance in arthritis, including TNF and IL-6, as well as to hypoxia, a known feature of the synovial environment[40]. The role of IFNγ in arthritis is less well understood. Studies performed more than 20 years ago implicated IFNγ in expression of the high-affinity Fc receptor FcγRI (CD64) on synovial fluids neutrophils[6]. Potential IFNγ sources in arthritis suggested by human and/or murine studies include CD4 T cells (IFNγ is the hallmark Th1 cytokine), CD8 T cells, NK cells, and NKT cells[41-46]. Experimental overexpression of IFNγ in the joint accelerates cartilage injury through upregulation of IgG Fc receptors and therefore enhanced susceptibility to immune complex injury[47]. However, IFNγ-deficient mice exhibit normal susceptibility to IgG-mediated K/BxN serum transfer arthritis, while IFNγ blockade or IFNγ receptor deficiency accelerates the onset and severity of collagen-induced arthritis[44, 48, 49]. Trials of recombinant IFNγ in rheumatoid arthritis found at best modest disease amelioration[50, 51]. These findings reflect the net impact of IFNγ on multiple lineages and remain compatible with the possibility that neutrophil exposure to IFNγ in arthritis is pro-inflammatory (for example, through upregulation of surface Fc receptors and HLA-DR), anti-inflammatory (for example, through up-regulation of the T cell inhibitor PD-L1), or both.

Comparing the transcriptional signature of human and murine neutrophils, we observed substantial overlap, including shared presence of an IFNγ signature in synovial fluid neutrophils, supporting the human relevance of extensive murine work defining the role of neutrophils in arthritis[11, 13-16]. This conclusion is important because performance of unambiguous human studies is complicated by the lack of neutrophil-specific therapeutics. Cross-species similarity is further echoed in murine neutrophil scRNAseq data, where differences between healthy and arthritic blood neutrophils were small whereas differences between arthritic blood and synovial neutrophils were large[19]. Whereas the neutrotime signature cannot be extrapolated directly to bulk RNAseq data, downregulation of early-neutrotime transcripts such as *LCN2, CAMP*, and *CD177* further supports the suggestion that human synovial fluid neutrophils – like their murine counterparts – skew similarly skew toward an aged phenotype reflecting prolonged survival in the inflamed joint environment[19].

Of particular interest is the marked cell-to-cell heterogeneity revealed by CyTOF. The clusters reported here reflect investigator-chosen analytical parameters and therefore are best regarded as one snapshot of this complex population rather than as discrete subsets. The data show that neutrophils within the inflamed joint differ phenotypically from each other as well as from those in blood. Dimensionality reduction by UMAP identified 2 broad populations: SFN1, resembling circulating neutrophils, and SFN2, a more abundant group often bearing markers associated with the IFNγ signature. We speculate that these populations reflect a chronological progression, with SFN1 representing recent arrivals that evolve in SFN2 cells with exposure to the inflamed synovial environment and time. This suggestion is consistent with greater expression within SFN2 of the maturity marker CD10 and the aging marker CXCR4, although SFN2 neutrophils remain internally diverse[52].

We applied both inflammatory stimuli and time to cultured healthy donor blood neutrophils. Indeed, two orthogonal signals were noted: IFNγ exposure upregulated hallmark SFN2 proteins such as HLA-DR, PD-L1, and CD64, while aging was required to yield the second key SFN2 marker, CXCR4 (interestingly partially suppressed by IFNγ). Further study will be required to confirm the parallels between these findings and the arthritis context, but the data nevertheless support the conceptual model that the neutrophil phenotypes observed in human synovial fluid represent an integration of inflammatory stimuli and aging in cells recruited in an ongoing manner to the inflamed joint.

Our work has several limitations. RNAseq studies employed bulk sorted neutrophils, enabling us to identify transcripts in depth but prohibiting us from calculating developmental trajectories. Future studies using scRNAseq will be important to define the ontological relationships among joint fluid neutrophils. CyTOF studies employed a set of antigens reflecting investigator choice; for technical reasons, not all antigens proved interpretable, and it is likely that many informative antigens were omitted. Our data do not detail epigenetic reprograming of neutrophils, and we did not characterize the function of the heterogeneous groups identified in arthritic synovial fluid. Despite these limitations, the results represent a uniquely granular examination of the transcriptional and surface/intracellular phenotype of human arthritic neutrophils, setting the stage for the next set of phenotypic and functional studies toward the ultimate goal of identifying targetable pathways for therapeutic neutrophil blockade in arthritis.

## Acknowledgements

We thank Adam T. Chicoine and the Brigham and Women’s Hospital cell sorting facility for cell sorting and Jüri Habicht for technical assistance.

## Patient consent for publication

Not required.

## Ethics approval

EDTA-anticoagulated blood from healthy donors was collected under IRB-approved protocols (BWH-2008P000427 and Heidelberg S-285/2015). Simultaneous blood and synovial fluid samples were obtained with written consent from patients with inflammatory arthritis who underwent diagnostic and/or therapeutic joint aspiration (BCH-P00005723 and BWH-2006P001068). Patients or the public were not involved in the design, or conduct, or reporting, or dissemination plans of our research.

## Data availability statement

All relevant data is included in the article and online supplemental information. RNA-sequencing data will be made publicly available upon publication.

