## Supplementary Information for "Aging and interferon gamma response drive the phenotype of neutrophils in the inflamed joint"

*** Correspondence** to R.G-B and P.A.N.:

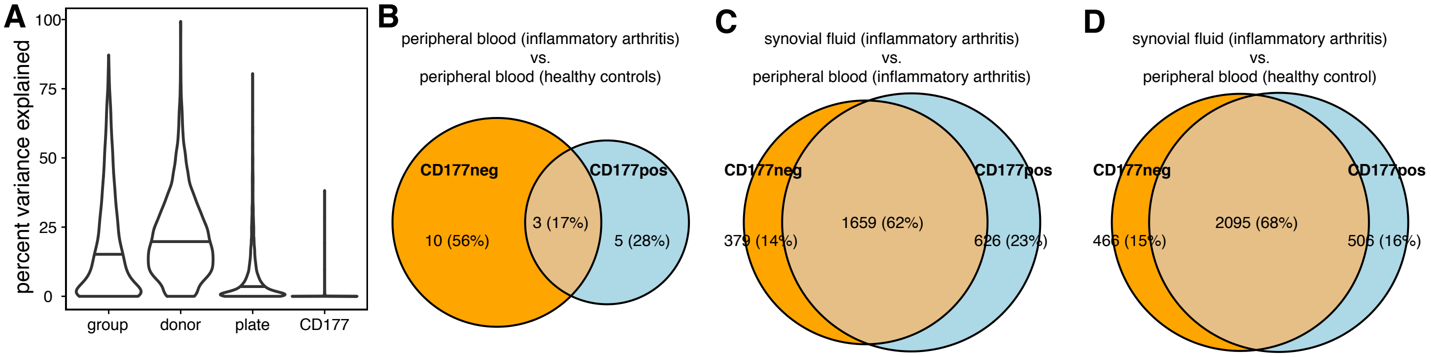


**Supplementary Figure 1. Overlap in the transcriptome of CD177-defined neutrophils.**
**(A)** Percentage of variance in gene expression explained by tissue, donor number, RNAseq batch (plate) and CD177 status.
**(B-D)** Overlap between differentially expressed genes in CD177neg and CD177pos neutrophils in different pairwise comparisons.

---

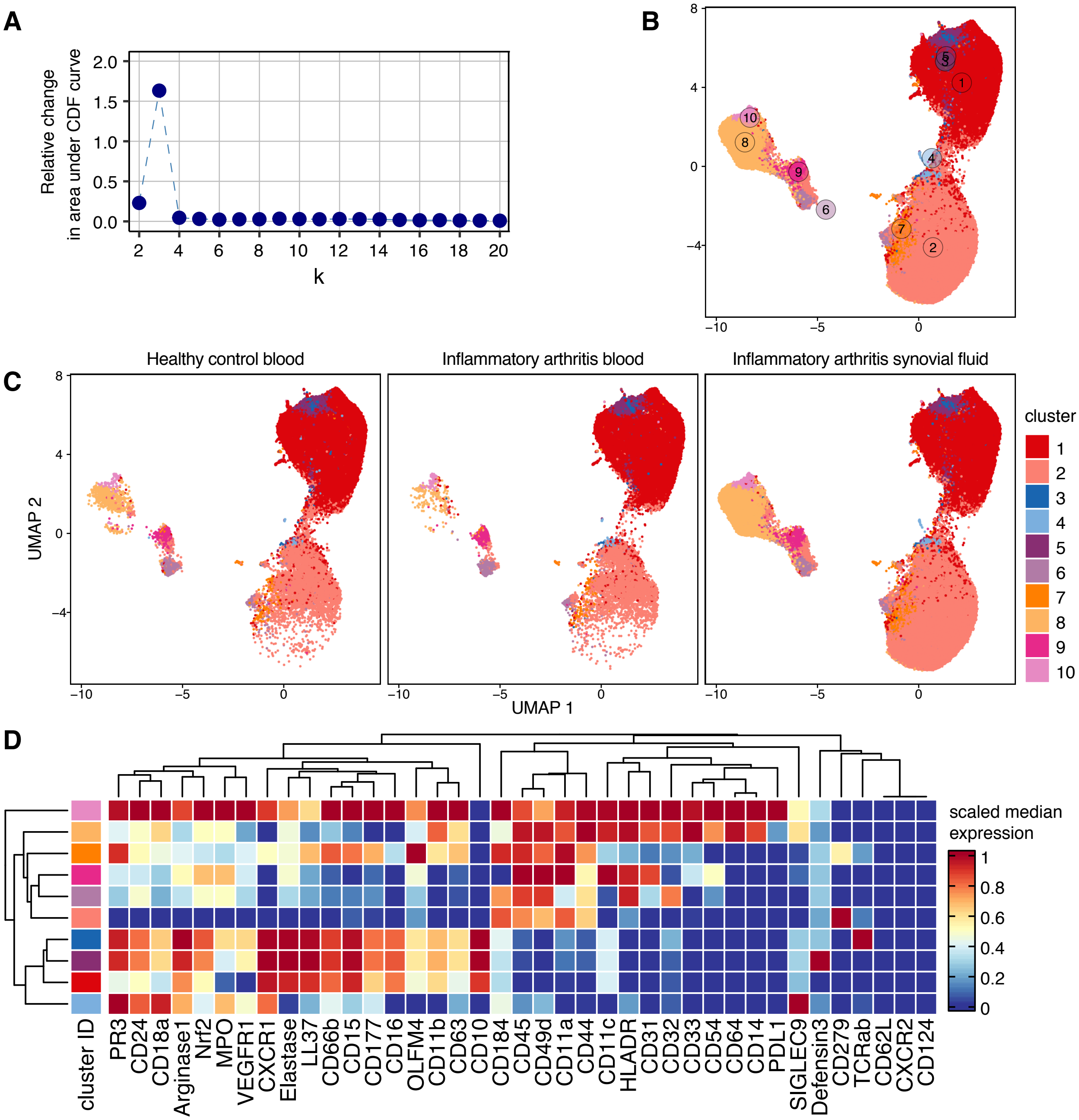


**Supplementary Figure 2. CyTOF gating strategy.**
**(A)** Delta area plot for increasing numbers of clusters. **(B)** UMAP analysis of all leukocytes, colored by k = 10 clusters. **(C)** UMAP analysis split by source tissue. **(D)** Marker expression of leukocytes across different clusters. Only cells belonging to clusters 1, 3 and 5 and with UMAP1 ≥ –1.5 and UMAP2 ≥ 0.5 were included in downstream analysis.

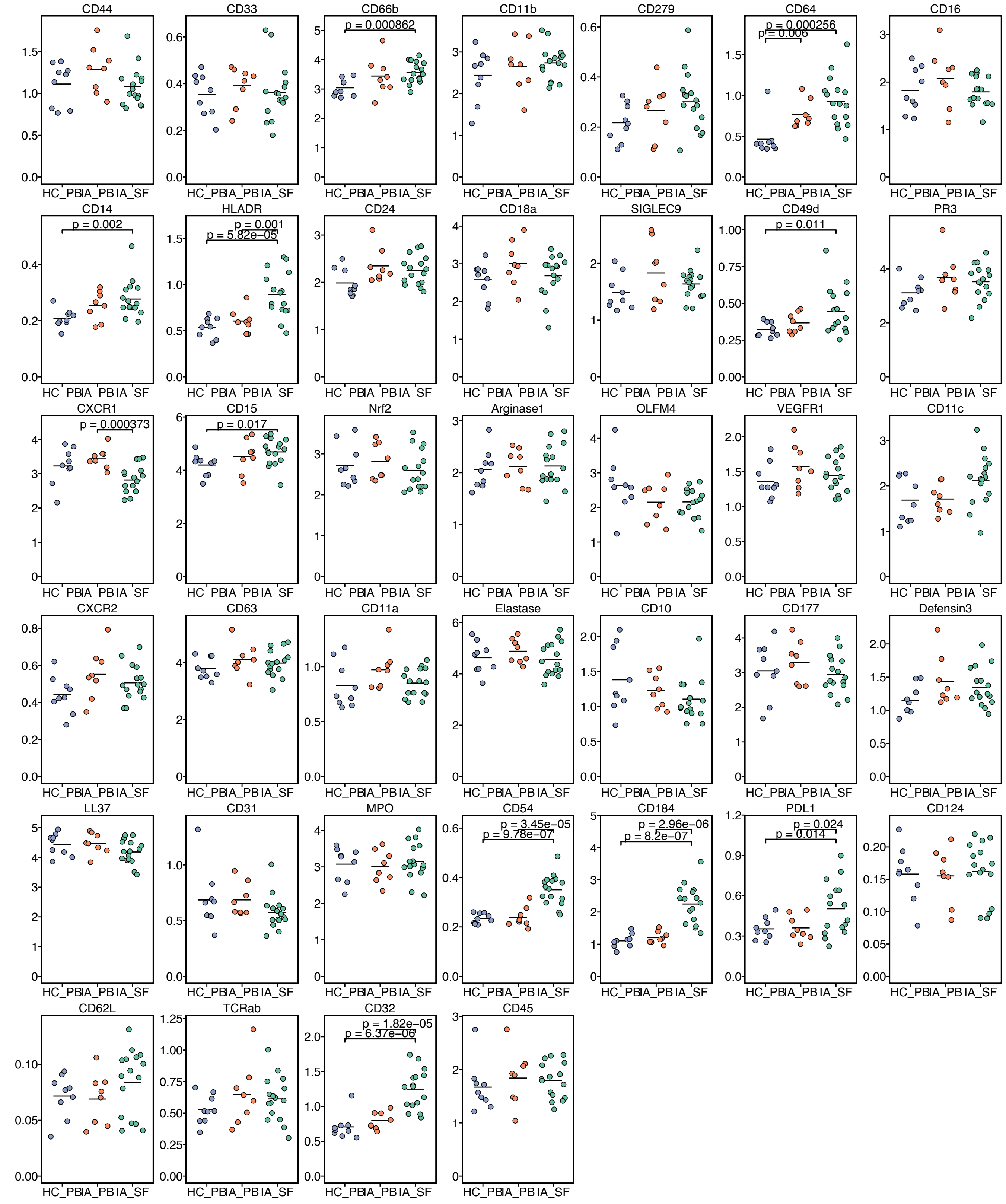


**Supplementary Figure 3. Differential expression analysis for all CyTOF markers between blood and synovial fluid.**
HC_PB = healthy control blood; IA_PB = inflammatory arthritis blood; IA_SF = inflammatory arthritis synovial fluid

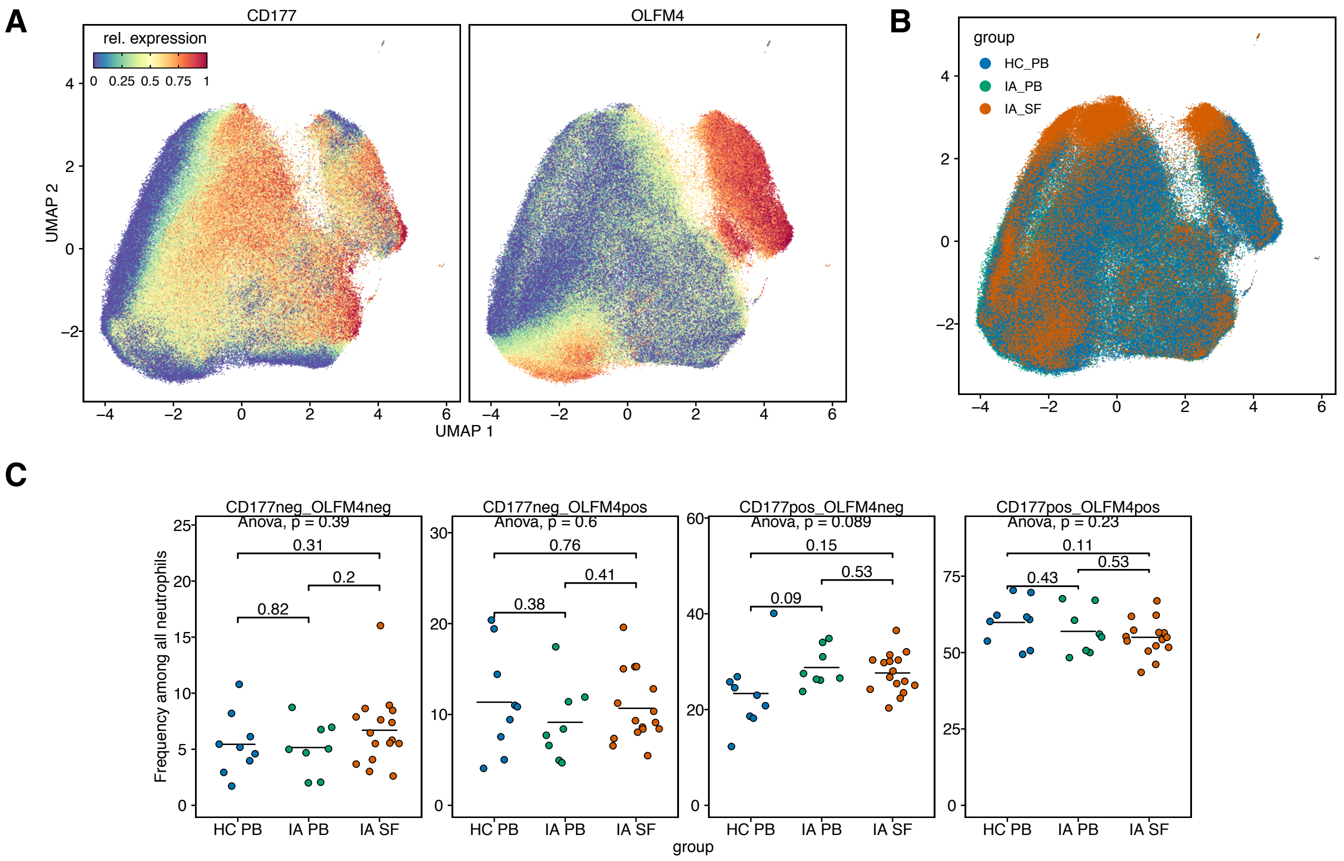


**Supplementary Figure 4. Single cell analysis of CyTOF data without exclusion of CD177 and OLFM4.**
**(A)** UMAP dimensionality reduction is dominated by the dichotomously expressed OLFM4 and CD177. **(B)** Healthy control blood, inflammatory arthritis blood and inflammatory arthritis synovial fluid cells are not enriched in specific areas of high/low CD177/OLFM4 expression. **(C)** No difference in frequency of OLFM4+/– or CD177+/– cells between the groups.
HC_PB = healthy control blood; IA_PB = inflammatory arthritis blood; IA_SF = inflammatory arthritis synovial fluid

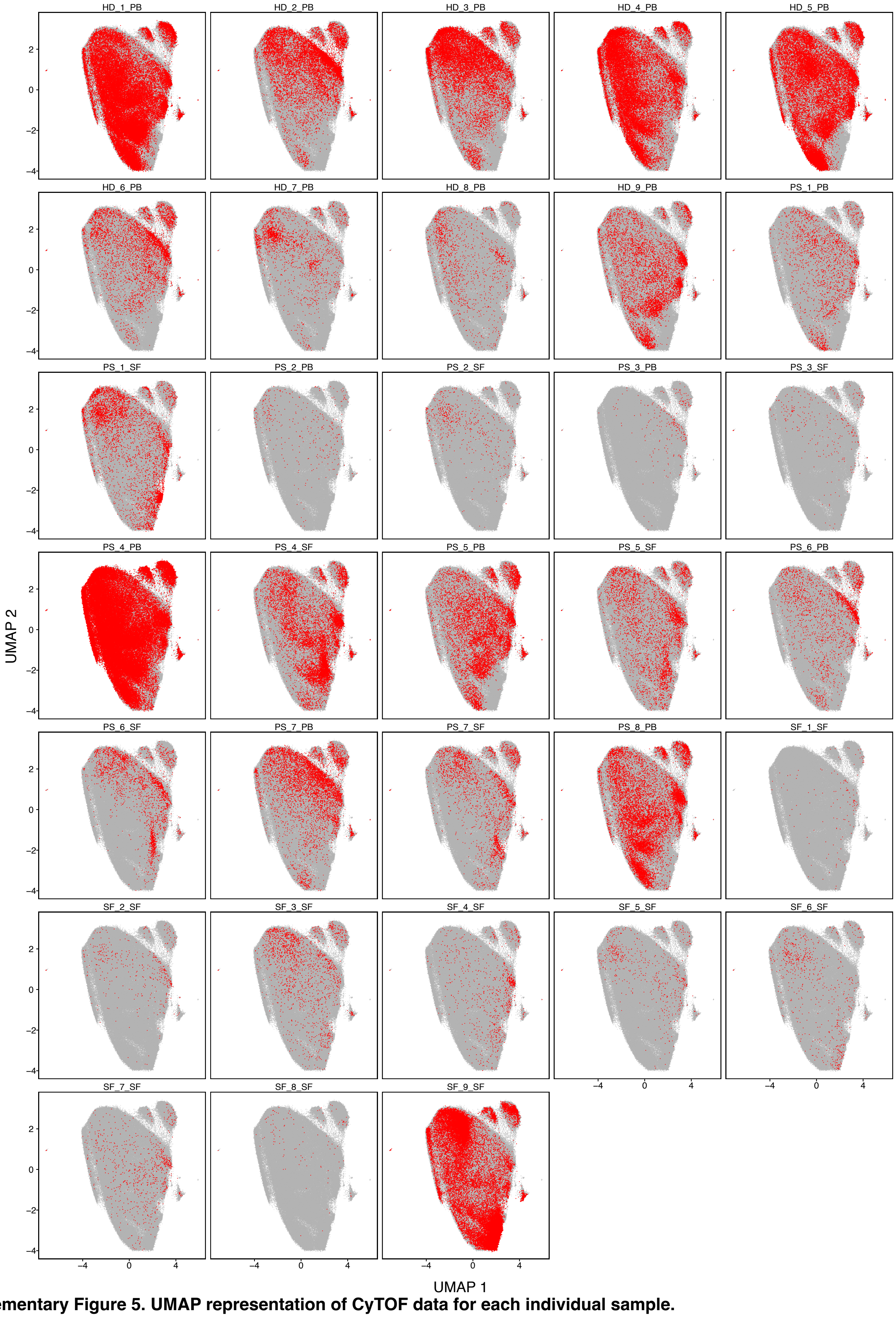


**Supplementary Figure 5. UMAP representation of CyTOF data for each individual sample.**
HD_PB = healthy donor blood; PS_PB/PS_SF = inflammatory arthritis paired blood/synovial fluid;
SF = inflammatory arthritis synovial fluid without matched blood

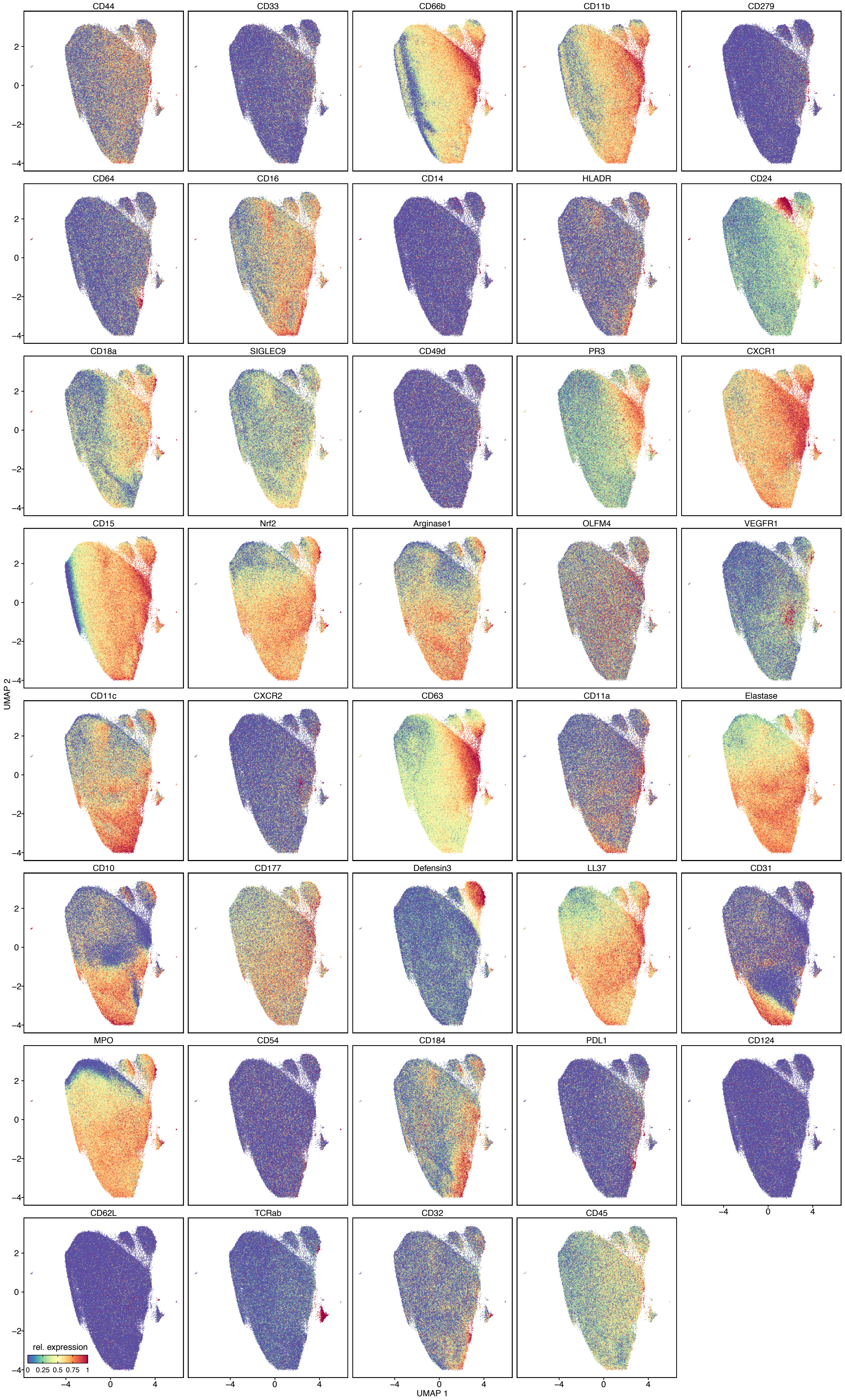


**Supplementary Figure 6. UMAP representation of all markers in the CyTOF data.**

### SUPPLEMENTARY METHODS

#### Research subjects

Transcriptomic studies were performed on peripheral blood from 15 healthy controls and on contemporaneous blood and synovial fluid from 16 patients with inflammatory arthritis including at least one active joint requiring therapeutic arthrocentesis. CyTOF was performed on blood from 9 healthy volunteer donors and in samples from 17 patients with inflammatory arthritis: 7 contemporaneous blood and synovial fluid, 1 blood only, and 9 synovial fluid only. Demographic characteristics of the donors are displayed in **Supplementary Table 1**.

**Supplementary Table 1. Description of the study population**

| | **RNAseq** | | **CyTOF** | |
|---|---|---|---|---|
| | **Healthy controls (N = 15)** | **Inflammatory arthritis (N = 16)** | **Healthy controls (N = 9)** | **Inflammatory arthritis (N = 17)** |
| **Age (years) (Mean, range) undisclosed (%)** | 32 (23–49)<br>6/15 undisclosed (40%) | 20.1 (2–73)<br>2/16 undisclosed (12%) | 34 (24–50) | 24 (4–67)<br>2/17 undisclosed (12%) |
| **Biological sex Male (%) Female (%) undisclosed (%)** | 10/15 female (67%)<br>5/15 male (33%) | 8/16 female (50%)<br>6/16 male (38%)<br>2/16 undisclosed (12%) | 3/9 female (33%)<br>6/9 male (67%) | 10/17 female (59%)<br>5/17 male (29%)<br>2/17 undisclosed (12%) |
| **Diagnosis** | | 2/16 rheumatoid arthritis (12%)<br>11/16 juvenile idiopathic arthritis (69%)<br>3/16 undifferentiated inflammatory arthritis (19%) | | 8/17 rheumatoid arthritis (47%)<br>8/17 juvenile idiopathic arthritis (47%)<br>1/17 undifferentiated inflammatory arthritis (6%) |

#### Neutrophil collection and cell isolation

EDTA-anticoagulated blood from healthy donors was collected under IRB-approved protocols (BWH-2008P000427 and Heidelberg S-285/2015). Simultaneous blood and synovial fluid samples were obtained with written consent from patients with inflammatory arthritis who underwent diagnostic and/or therapeutic joint aspiration (BCH-P00005723 and BWH-2006P001068).

Neutrophils were isolated from fresh blood by negative selection using the EasySep™ Direct Human Neutrophil Isolation Kit (STEMCELL #19666), routinely yielding a purity > 90% as determined by flow cytometry. For synovial fluid, in case of visible erythrocyte contamination, up to 5 ml were mixed with 30 ml ammonium-chloride based RBC lysis buffer for 5 minutes and then topped up to 50 ml with RPMI containing 1% FBS. Cells were pelleted at 400 × g for 5 min and used fresh for cell sorting, as well as cryopreserved in Cryostor CS10 (frozen at -80 ºC and stored in liquid nitrogen) until CyTOF.

#### Cell sorting for RNAseq

Pelleted cells were resuspended in antibody staining cocktail containing conjugated antibodies against CD15 (AF488), CD66b (AF647) and CD177 (BV421) at 1:100 (all antibodies from Biolegend). Cells were stained for 15 minutes on ice, followed by washing with RPMI 1% FBS. Cells were sorted on a BD Aria Fusion cell sorter into 1.5 ml Eppendorf tubes containing 350 µl RLT buffer and stored at -80 ºC awaiting RNA extraction. 30,000 $CD177^{pos}$ and 30,000 $CD177^{neg}$ neutrophils were sorted from each sample. Excess cells were sorted, pelleted and resuspended in 1 ml RLT buffer per million neutrophils for RNA extraction. CD177 is the only known surface molecule to segregate neutrophils into discrete populations independent of developmental stage or activation state, in most individuals based on bimodal expression[1, 2]. We thus sorted CD15+ CD66b+ neutrophils into $CD177^{pos}$ and $CD177^{neg}$ groups, reasoning that each group could first be analyzed separately and later merged within each donor. OLFM4 is another neutrophil protein with bimodal expression pattern, but the intracellular localization prohibited sorting based on this marker[3].

#### RNA extraction, SmartSeq2 library preparation and sequencing

RNA was extracted from lysed neutrophils using the Qiagen RNeasy micro kit following manufacturer instructions. RNA sequencing was performed at the Broad Institute under the Smartseq2 protocol[4]. Samples were sequenced on an Illumina NextSeq500with 25bp paired-end reads and an average sequencing depth of $6 \times 10^6$ reads per sample.

#### Analysis of RNAseq data

Short sequence reads were aligned to the reference genome (GRCh38) and Ensembl [5] transcriptome using STAR aligner[6]. The uniquely aligned reads were used to quantify gene expression levels for all Ensembl genes. Genes with low expression (less than 10 reads in at least three samples in at least one group) were removed. Data was analyzed in limma [7] using voom normalization [8] and the expression of each gene was modeled as a combination of donor- and group-specific effects. Genes were tested for differential expression with a false discovery rate (FDR) 0.05 as indicated. Gene Set Enrichment Analysis was performed using fgsea [9] on T score pre-ranked gene lists.

CD177 status accounted for less than 1% of variance compared with inter-individual (~20%) and tissue (blood vs. synovial fluid, ~15%) effects **(Supplementary Figure 1A)**. Gene expression changes across conditions were similar in CD177-defined populations **(Supplementary Figure 1B)**. $CD177^{pos}$ and $CD177^{neg}$ neutrophils from each donor were thus merged again to create bulk profiles. Differential expression testing was performed separately for $CD177^{neg}$ and $CD177^{pos}$ neutrophils and only genes displaying significant changes in both comparisons and in the same direction were considered in downstream analysis. For example, of 2,038 differentially expressed genes in $CD177^{neg}$ neutrophils between synovial fluid and peripheral blood of patients with inflammatory arthritis, 1,659 (81%) were also differentially expressed in CD177$^{pos}$ neutrophils **(Supplementary Figure 1B)**. Correspondingly, in CD177$^{pos}$ neutrophils, of 2,285 differentially expressed genes, 1,659 (73%) were also differentially expressed in CD177$^{neg}$ neutrophils.

### Cytometry by Time of Flight (CyTOF)

CyTOF was employed in a single batch to characterize the expression of 39 surface and intracellular proteins using a custom antibody panel **(Supplementary Table 2)**. Blood and synovial fluid samples were washed and stained with metal-conjugated monoclonal antibodies. For all stains, viability staining by cisplatin was added to plated cells for 10 min before staining. After washing and centrifugation, Fc-Block reagent was added for 10 min before adding cell-surface CyTOF antibody staining cocktails. Cells were stained for 30 minutes, then washed once in CyTOF staining buffer (CSB, calcium/magnesium-free PBS, 0.2% BSA, 0.05% sodium azide). Cells were fixed with 1.6% paraformaldehyde diluted in PBS (PFA) for 10 minutes, washed by centrifugation, then permeabilized with eBioscience FOXP3/Transcription Factor Staining Buffer (Life Technologies # 00-5523-00) for sample barcoding with palladium reagents as described [10]. After barcoding, cells were washed by centrifugation, combined into a single tube, fixed with Fix/Perm buffer, then washed once in permeabilization buffer. The intracellular CyTOF antibody staining cocktail (antibodies against PR3, Nrf2, Arginase 1, OLFM4, Elastase, Defensin 3, LL-37 and MPO) was diluted in permeabilization buffer to stain cells for 1 hour. Cells were washed in CSB by centrifugation, then fixed overnight in 1.6% PFA. Prior to acquisition, iridium-intercalator solution (MaxPar Intercalator-Ir 500 mM; Fluidigm Sciences) was added to cells in PBS. After washing by centrifugation, cells were diluted in Cell Acquisition Solution (CAS, Fluidigm Sciences) at a concentration of 7.5 x $10^5$ cells/ml with addition of EQ calibration beads (EQ Four Element Calibration Beads; Fluidigm Sciences) to normalize metal intensity signals using the manufacturer's protocol. Cells were analyzed on a Helios mass cytometer (Fluidigm Sciences).

**Supplementary Table 2 – Mass cytometry panel**

| Antigen | Clone | Metal | Dilution | Antigen | Clone | Metal | Dilution |
|---|---|---|---|---|---|---|---|
| **CD44** | IM7 | 113In | 1:100 | **CD11c** | Bu15 | 159Tb | 1:100 |
| **CD33** | WM53 | 115In | 1:100 | **CXCR2** | 5E8/CXCR2 | 160Gd | 1:100 |
| **CD66b** | G10F5 | 141Pr | 1:100 | **CD63** | H5C6 | 161Dy | 1:100 |
| **CD11b** | M1/70 | 142Nd | 1:100 | **CD11a** | HI111 | 162Dy | 1:100 |
| **CD279/PD1** | EH12.2H7 | 143Nd | 1:100 | **Elastase *** | 950317 | 163Dy | 1:100 |
| **CD64** | 10,1 | 144Nd | 1:100 | **CD10** | HI10A | 164Dy | 1:100 |
| **CD16** | 3G8 | 145Nd | 1:100 | **CD177** | MEM-166 | 165Ho | 1:100 |
| **CD14** | M5E2 | 146Nd | 1:100 | **Defensin 3 *** | Polyclonal | 166Er | 1:100 |
| **HLA-DR** | L243 | 147Sm | 1:100 | **LL-37 (Cathelicidin) *** | H7 | 167Er | 1:800 |
| **CD24** | ML5 | 148Nd | 1:100 | **CD31** | WM59 | 168Er | 1:100 |
| **CD18 (a)** | MEM-148 | 149Sm | 1:50 | **MPO *** | 392105 | 169Tm | 1:50 |

| | | | | | | | |
|---|---|---|---|---|---|---|---|
| **SIGLEC9 (CD329)** | K8 | 150Nd | 1:100 | **ICAM (CD54)** | HA58 | 170Er | 1:100 |
| **CD49d** | 9F10 | 151Eu | 1:100 | **CD184/CXCR4** | 12G5 | 171Yb | 1:100 |
| **PR3 *** | 684022 | 152Sm | 1:800 | **PDL1** | 29E.2A3 | 172Yb | 1:100 |
| **CXCR1** | 8F1 | 153Eu | 1:50 | **CD124** | G077F6 | 173Yb | 1:100 |
| **CD15** | MC-480 | 154Sm | 1:100 | **CD62L** | DREG-56 | 174Yb | 1:100 |
| **Nrf2 *** | 383727 | 155Gd | 1:50 | **TCR apha/beta** | IP26 | 175Lu | 1:100 |
| **Arginase 1 *** | 14D2C43 | 156Gd | 1:100 | **CD32** | FUN-2 | 176Yb | 1:100 |
| **OLFM-4 *** | 806305 | 157Gd | 1:50 | **CD45** | HI30 | 209Bi | 1:100 |
| **VEGFR1** | Polyclonal | 158Gd | 1:100 | | | | |

* denotes intracellular staining

### Analysis of CyTOF data

Data analysis was conducted using FlowJo version 10.7.1 and R. During initial quality control, contaminating leukocytes were removed **(Supplementary Figure 2)**. We retained 280,174 single neutrophils from a total of 33 samples in our dataset. These comprised 119,354 neutrophils from healthy control blood, 94,771 neutrophils from blood of patients with inflammatory arthritis, and 66,049 neutrophils from inflamed synovial fluid. We extracted mean signal intensities for each channel per sample and calculated Spearman correlation coefficients between samples as with the RNAseq data. Using these mean expression values, fold changes were calculated and $\log_2$ transformed. For marker expression between groups, multiple t-tests were performed and p-values adjusted using Holm correction. For marker-to-marker correlation analysis, intensity on the bulk level was defined as the mean intensity of each marker from all the cells in each sample and on the single cell level, each cell was considered separately. For analysis on the single-cell level, UMAP dimensionality reduction was performed with standard settings, using all antigens (Supplementary Figures 2 and 4) or all antigens minus OLFM4, CD177 and PR3 (all other Figures) as input. Neutrophils were intentionally overclustered into k = 20 clusters. Abundance of populations between groups was tested using ANOVA, followed by independent t-tests.

### Neutrophil stimulation

For functional experiments, cells were isolated from heparin-anticoagulated blood by density gradient centrifugation as described previously[11]. Briefly, 30 ml of blood were layered on top of 20 ml PolymorphPrep (Progen #1114683) and centrifuged for 35 min at 535 g at room temperature (23 °C). Neutrophils were recovered and contaminating red blood cells were removed by hypotonic lysis. The neutrophils were subsequently cultured at $10^6$ cells/ml for up to 48 h in RPMI 1640 (Gibco #21875-034) supplemented with 10% heat-inactivated FBS (PAN Biotech #3302/P101102) and 1% GlutaMAX (Gibco #35050-061) in a humidified atmosphere at 37°C and 5% $CO_2$. For cytokine stimulation, IFNγ (BioLegend #570208) was added to the medium at 10 ng/ml. Cells cultured in medium for the indicated durations without the addition of cytokines served as controls. At the indicated timepoints, $10^6$ cells were harvested and stained for live/dead discrimination and an antibody panel **(Supplementary Table 3)** in a total volume of 50 µl for 20 and 25 min, respectively. Stained cells were resuspended in FACS buffer containing 2% FBS, 5 mM EDTA and 0.1% sodium azide in PBS and 50,000 events per sample were recorded on a BD LSRII flow cytometer. FCS files were analyzed and gated with FlowJo (v. 10.8.0.) and live CD66b+ cells as well as the corresponding median fluorescence intensities were exported for further analysis.

#### Supplementary Table 3 – Flow cytometry panel

| Marker | Channel | Clone | Vendor | Catalog # | dilution |
|---|---|---|---|---|---|
| **LIVE/DEAD** | Pacific Orange | N/A | BioLegend | 423103 | 1:300 |
| **HLA-DR** | AF700 | L243 | BioLegend | 307626 | 1:100 |
| **PD-L1** | APC | MIH1 | BD BioScience | 563741 | 1:20 |
| **CD66b** | APC-Cy7 | G10F5 | BioLegend | 305126 | 1:100 |
| **CD32** | FITC | FUN-2 | BioLegend | 303204 | 1:100 |
| **CD64** | PE | 10.1 | BioLegend | 305007 | 1:100 |
| **CXCR4** | BV421 | 12G5 | BioLegend | 306518 | 1:100 |
| **ICAM-1** | PE-Cy5 | HA58 (RUO) | BD BioSciences | 555512 | 1:25 |

#### Flow cytometry analysis

FCS files were concatenated and analyzed using the CATALYST pipeline (v. 1.16.2) in R (v. 4.1.0). Fluorescence values were arcsinh transformed with manually determined cofactors [12] and subsequently clustered by FlowSOM-Clustering followed by ConsensusMetaClustering. Dimensionality reduction was performed using the DiffusionMap algorithm within the CATALYST package with standard settings on a random subset of 5000 cells per sample for better visualization. Plots were generated using ggplot2 (v. 3.3.5; (4)) and CATALYST. Median fluorescence values were assembled and plotted using the ggplot2 package in R.
